# Simulated microgravity induces aging phenotypes in human organoids

**DOI:** 10.64898/2026.09.30.755771

**Authors:** Wei-Chieh Mu, Jordan Baechle, Kevin Schneider, Matias Fuentealba, Fei Wu, Max Manwaring-Mueller, Nathan Bracey, Alexander Chebykin, Abhayjit Saini, Jiwoon Park, JangKeun Kim, Daniel A. Winer, Nazish Sayed, Chikage Anna Noishiki, Alan I. Gale, Christopher E. Mason, Mark M. Davis, David Furman

## Abstract

Simulated microgravity provides a controlled perturbation for testing how loss of mechanical load affects human tissues. After 24 hours in rotating-wall vessels, neural, cardiac and tonsil organoids developed tissue-specific transcriptional responses that overlapped published spaceflight datasets, accompanied by higher transcriptomic-age estimates in neural and cardiac models, altered cardiac-organoid contraction dynamics and modified tonsil-organoid antiviral gene-expression signatures. These findings establish a multi-organoid framework for studying acute responses to gravitational unloading.

## Main

Spaceflight perturbs neural, cardiovascular and immune physiology, and several responses resemble molecular features associated with aging^1-4^. The Space Omics and Medical Atlas and related studies have begun to resolve these effects at the molecular level, including mitochondrial and immune responses^5,6^, but differences among missions, exposure conditions, and experimental platforms complicate mechanistic interpretation. Whether these molecular responses translate into tissue-specific functional changes remains unclear, partly because controlled human models permitting parallel molecular and functional analysis are limited. To address this gap, we used three human organoid systems to test whether a common, short-duration simulated-microgravity exposure could recapitulate components of spaceflight-associated transcriptional changes, engage aging- and disease-related pathways, and alter tissue-level functional responses. We generated forebrain and cardiac organoids from human induced pluripotent stem cells (iPSC) and tonsil organoids from primary human lymphoid tissue, exposed them for 24 hours to simulated microgravity in rotating-wall vessels^7^, and analyzed their transcriptional and functional responses (Fig. 1a). Matched 1g controls enabled comparisons across organoid types and with human tissues and in vitro low-Earth orbit datasets.

**Figure 1.**
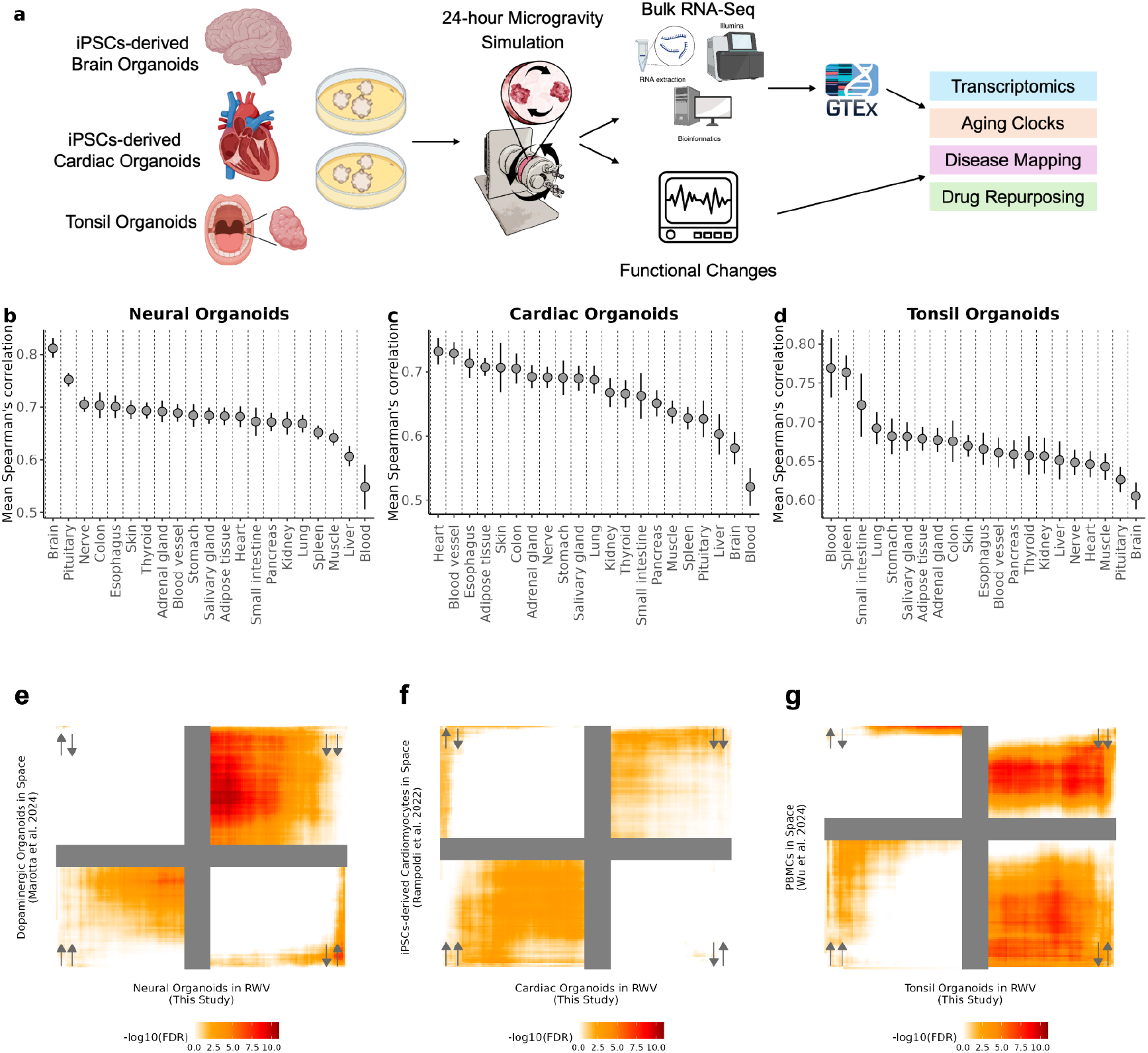
Tissue identity and correspondence with spaceflight-associated transcriptional signatures. a, Experimental workflow for neural, cardiac and tonsil organoids exposed for 24 hours to 1g or simulated microgravity (μG). b–d, Spearman correlations between 1g organoid transcriptomes and GTEx tissues. e–g, RRHO maps comparing ranked differential-expression profiles from rotating-wall-vessel organoids (x-axis) with published spaceflight datasets for neural, cardiac and immune cell signatures (y-axis), respectively. Color intensity represents the -log_10_ of the Benjamini-Hochberg adjusted hypergeometric overlap P value. Arrows indicate ranked-list directionality, where the left arrow represents the y-axis signature and the right arrow represents the x-axis signature.

Before evaluating the perturbation, we benchmarked the organoid transcriptomes against human tissues in the Genotype–Tissue Expression (GTEx) resource by calculating their Spearman rank correlation coefficients across 21 distinct GTEx tissues (Fig. 1b-d)^8^. Forebrain organoids correlated most strongly with the brain tissues (Fig. 1b), cardiac organoids with heart tissues (Fig. 1c), and tonsil organoids with blood and spleen (Fig. 1d). These comparisons support broad tissue identity, albeit without complete cellular composition, epigenome profiles, or tissue maturity. Rank–rank hypergeometric overlap (RRHO) analysis identified correspondence between simulated-microgravity-associated gene expression changes and published spaceflight signatures from space-flown dopaminergic neural organoids^9^ (Fig. 1e), space-flown iPSC-derived cardiomyocytes^10^ (Fig. 1f) and astronaut peripheral blood mononuclear cell (PBMC) signatures analyzed postflight versus preflight^11^ (Fig. 1g). The neural comparison showed predominantly concordant down-down gene expression overlap. The cardiac comparison showed predominantly concordant up-up gene expression overlap. The tonsil comparison identified prominent concordant down-down overlap and discordant overlap between genes upregulated in microgravity-exposed tonsil organoids and genes downregulated in astronaut PBMCs after spaceflight. Thus, the in vitro perturbation recovered components of independently observed spaceflight-associated transcriptional responses across three tissue contexts.

We next examined whether microgravity-associated transcriptional changes were enriched for gene sets associated with the twelve hallmarks of aging^12^. The three organoid types exhibited partially overlapping patterns. Neural organoids showed prominent enrichment signals involving deregulated nutrient sensing, cellular senescence, loss of proteostasis, and mitochondrial dysfunction. Cardiac organoids showed prominent signals involving cellular senescence, loss of proteostasis, mitochondrial dysfunction, altered intercellular communication and chronic inflammation. Tonsil organoids showed prominent signals involving cellular senescence, chronic inflammation, loss of proteostasis, altered intercellular communication and epigenetic alterations (Fig. 2a–c).

**Figure 2.**
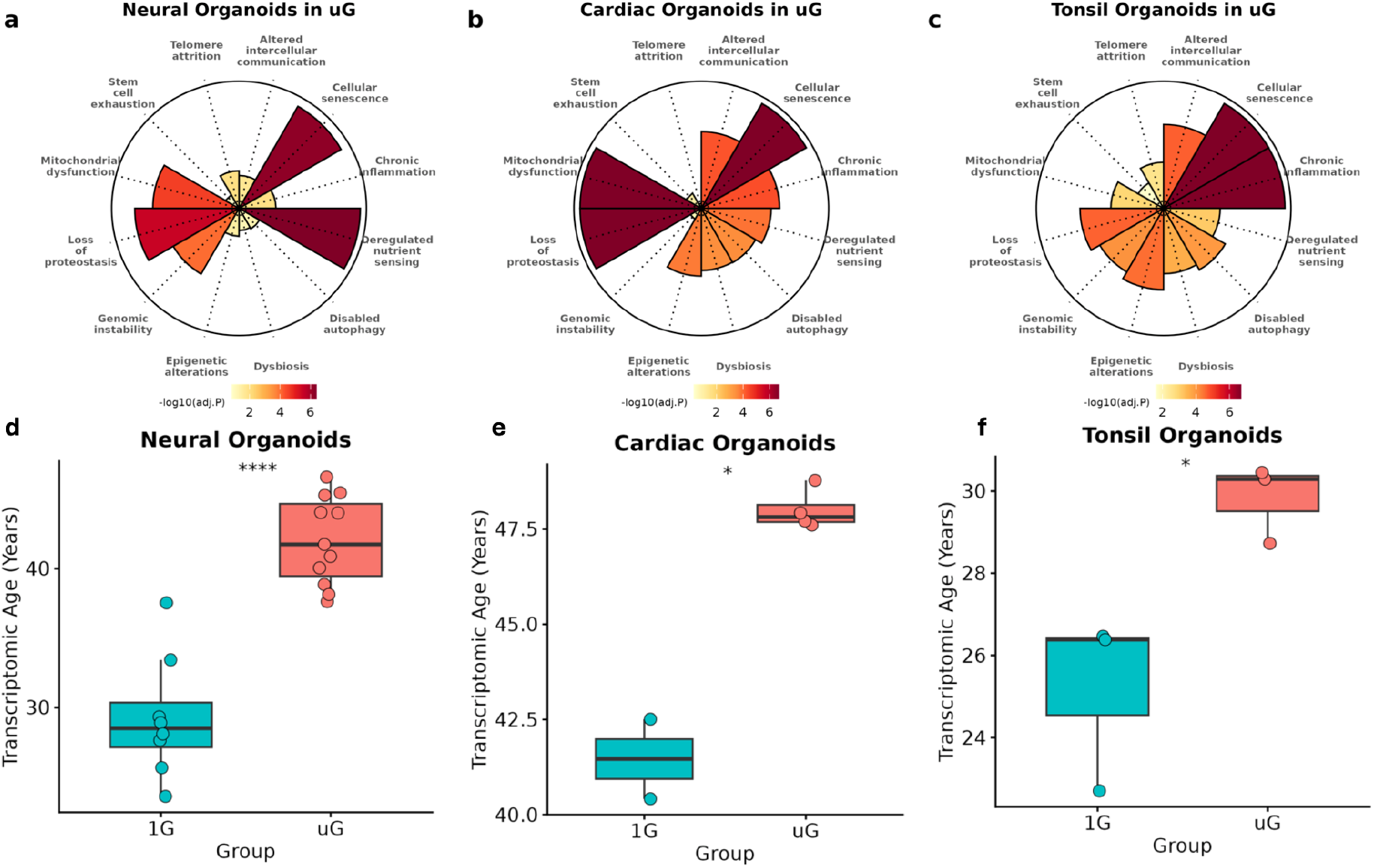
Aging-hallmark transcriptional signatures and predicted transcriptomic age. a–c, Enrichment of gene sets representing the twelve hallmarks of aging after 24 hours of simulated microgravity in neural, cardiac and tonsil organoids. Wedge length and color encode -log10(adjusted P value). d–f, Transcriptomic-age estimates from tissue-specific GTEx-trained models for neural and cardiac organoids and transcriptomic aging signature score for tonsil organoids maintained at 1g or exposed to simulated microgravity. Points represent individual samples; boxes summarize the sample distributions. Significance annotations are reproduced from the source analysis (****P < 0.0001; *P < 0.05; ns, not significant).

Tissue-specific transcriptomic aging clocks trained using GTEx brain-cortex and heart-left-ventricle reference data assigned higher age estimates to microgravity-exposed neural and cardiac organoids than to their 1g controls, with median shifts of approximately ten years for cortex-like tissue and five to seven years for ventricular myocardium (Fig. 2d, e). The transcriptomic aging signature derived from tonsil organoids made from dissociated tonsils extracted from old and young individuals significantly increased in tonsil organoids exposed to microgravity compared to their 1g controls, with median shifts of approximately 4 years (Fig. 2f).

These results represent substantial shifts in model-predicted transcriptomic age in neural, cardiac, and tonsil organoids, consistent with an accelerated aging-associated transcriptional profile.

Disease-gene-set analysis using DisGeNET^13^ linked the neural response to categories that included Parkinson’s disease, Alzheimer’s disease and Lewy Body disease (Fig. 3a). Several genes associated with Parkinson’s disease, neuronal identity, mitochondrial function, and cellular stress, including *PRKN* and *SNCB* showed reduced expression under simulated microgravity (Fig. 3b, c). *PRKN* encodes parkin, a Parkinson’s disease-associated ubiquitin ligase involved in mitochondrial quality control^14^. *SNCB* encodes β-synuclein, a synuclein-family protein implicated in synaptic function that, in experimental systems, has been reported to inhibit aggregation of *α*-synuclein, a major protein component of Lewy bodies in Parkinson’s disease^15^. Validation with qPCR showed reduced *SNCB* expression under microgravity (*P* = 0.025); *PRKN* showed a nonsignificant downward trend (*P* = 0.092) (Fig. 3c). Together, these findings suggest that microgravity may affect the expression of genes involved in synuclein biology and mitochondrial quality control, processes relevant to Parkinson’s disease.

**Figure 3.**
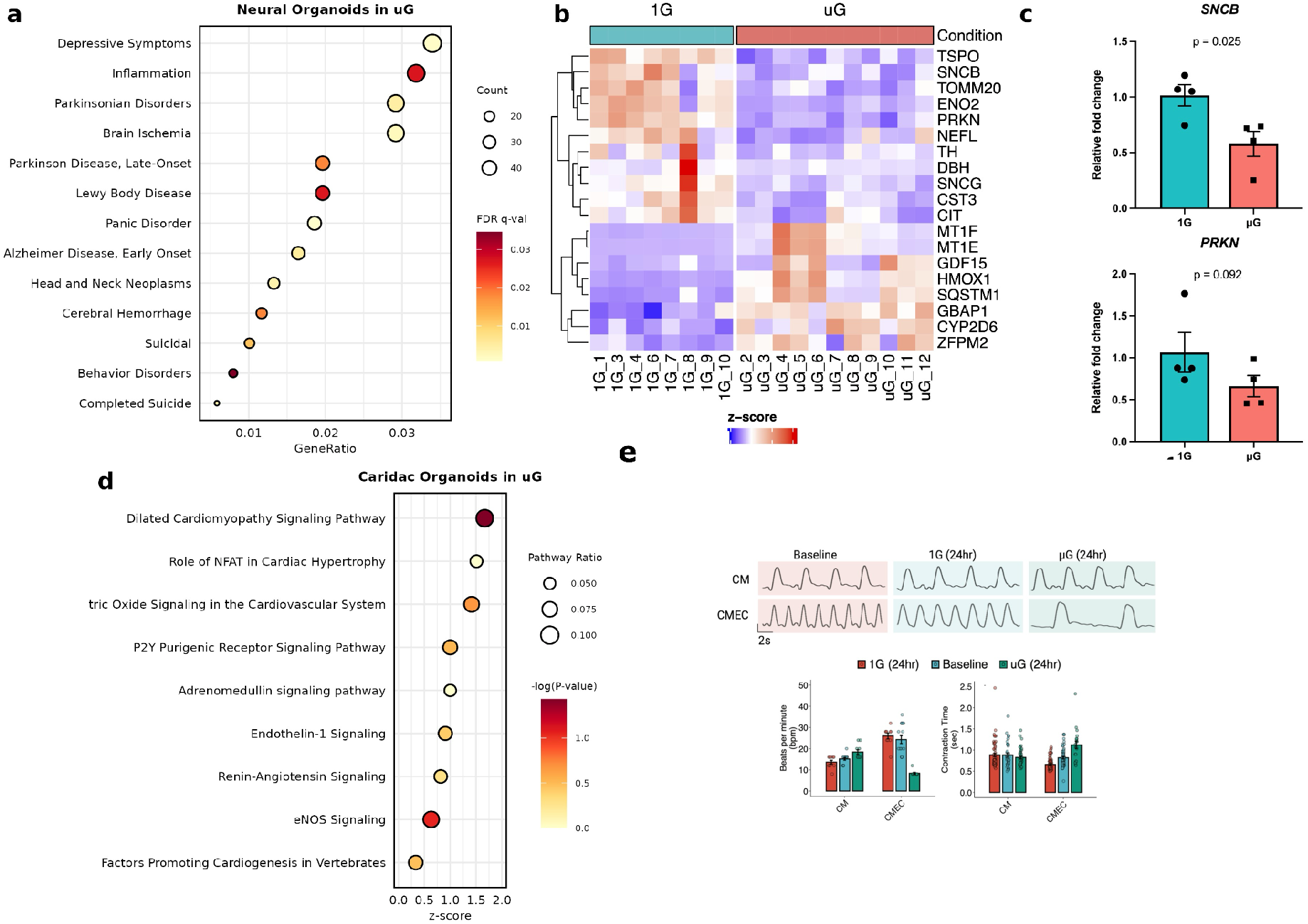
Neural disease associations and cardiac functional effects. a, DisGeNET overrepresentation analysis of differentially expressed genes in neural organoids after simulated microgravity; point size denotes gene count, and color denotes FDR q-value (lighter colors indicating greater significance). b, Row-standardized expression (z-scores) of selected Parkinson’s disease-associated genes. c, Relative SNCB and PRKN expression measured by qPCR. d, Cardiac pathway analysis; point size denotes pathway ratio, color denotes -log10(P value) (darker colors indicating greater significance) and horizontal position denotes activation z-score. e, Representative contraction traces and summary measurements for cardiomyocyte-only (CM) and cardiomyocyte–endothelial co-culture (CMEC) organoids at baseline and after 24 hours at 1g or μG.

Cardiac organoids showed enrichment of pathways related to dilated cardiomyopathy, cardiac remodeling, and cardiovascular signaling (Fig. 3d). Consistent with a functional cardiac phenotype, high-resolution video-based motion tracking showed a reduction in spontaneous beat frequency and a prolongation of contraction-relaxation cycles after 24 hours of simulated microgravity, most prominently in organoids containing both cardiomyocytes and endothelial cells (Fig. 3e).

Tonsil organoids provide a human model of adaptive immune responsiveness^16^. Live attenuated influenza vaccine (LAIV) elicited a broad transcriptional response, including upregulation of multiple antiviral and interferon-stimulated genes, demonstrating functional responsiveness to immune stimulation (Fig. 4a). Disease-gene-set analysis of the microgravity-associated response identified enrichment for categories related to infection, inflammation and immune-system regulation (Fig. 4b). RRHO analysis of LAIV-versus-unstimulated and microgravity-versus-1g transcriptional signatures suggests a predominantly discordant relationship between the two ranked gene-expression profiles (Fig. 4c). Consistent with this pattern, selected antiviral-response genes showed broadly positive log_2_ Fold Changes (log_2_FC) in the LAIV-versus-unstimulated contrast, and predominantly negative changes in the microgravity-versus-1g contrast (Fig. 4d). Together, these findings indicate an inverse relationship between the microgravity-and LAIV-associated transcriptional signatures, including expression of several antiviral and interferon-associated genes under microgravity. Testing whether microgravity dampens the response to LAIV will require a direct comparison of the LAIV response under 1g and microgravity conditions using a vaccine-by-gravity interaction model.

**Figure 4.**
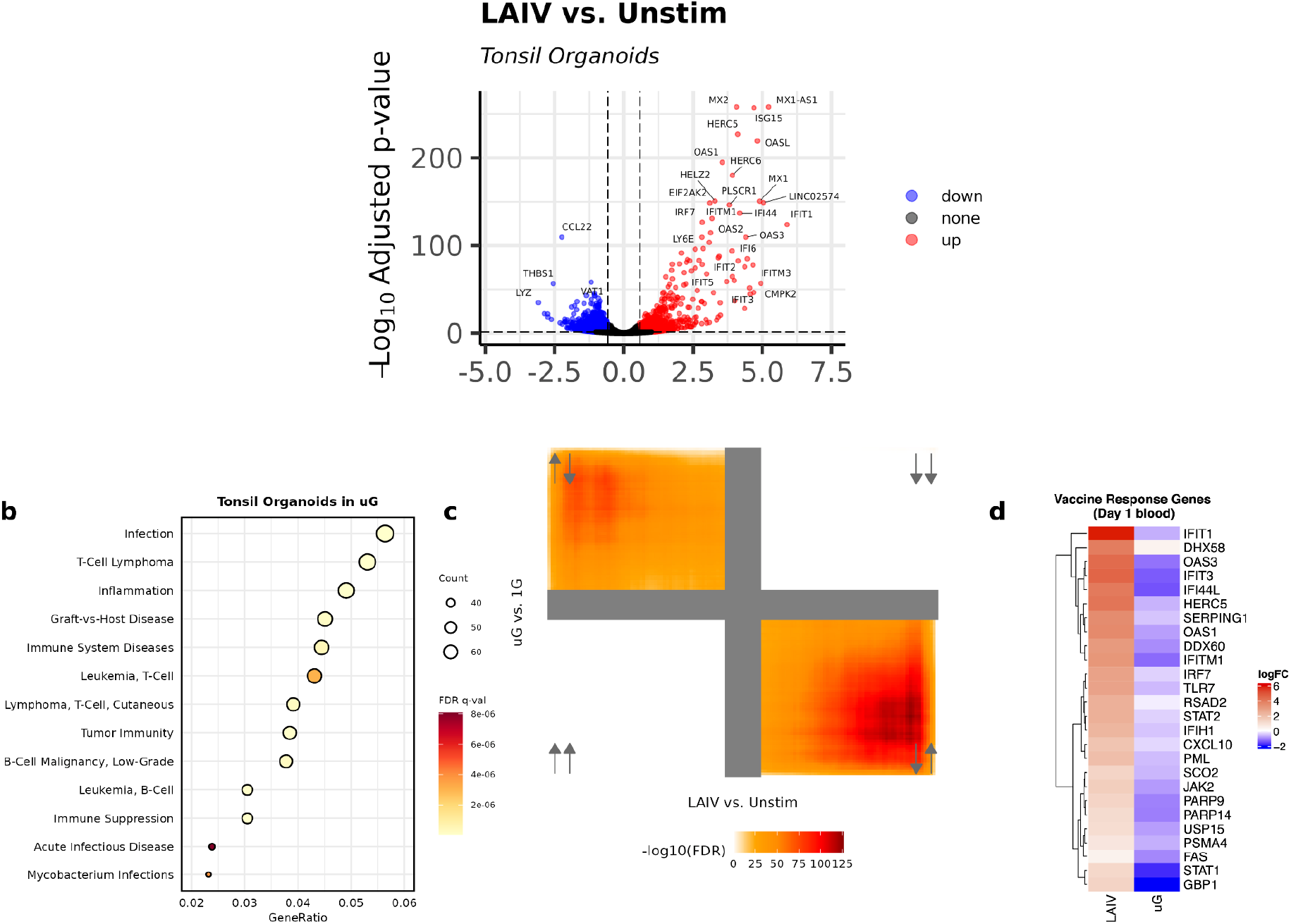
LAIV- and microgravity-associated transcriptional responses in tonsil organoids. a, Volcano plot comparing live attenuated influenza vaccine (LAIV)-stimulated and unstimulated tonsil organoids; color indicates the direction of differential expression. b, DisGeNET overrepresentation analysis after simulated microgravity; point size denotes gene count, and color denotes FDR q-value (lighter colors indicating greater significance). c, RRHO comparison of the LAIV-versus-unstimulated signature (x-axis) and the microgravity-versus-1g signature (y-axis). Color represents -log_10_ of the Benjamini-Hochberg adjusted hypergeometric overlap P value. d, log_2_FC for selected antiviral-response genes in the LAIV-versus-unstimulated and μG-versus-1g contrasts.

In sum, these results show that even 24 hours of simulated microgravity induces tissue-dependent transcriptional remodeling in human organoids and that components of these responses correspond to published spaceflight-associated signatures. By enabling matched molecular and functional measurements across several human tissue contexts, this platform provides a tractable, ground-based approach for prioritizing tissue-specific mechanisms for subsequent validation in spaceflight studies. Higher transcriptomic-age estimates in neural and cardiac organoids, altered cardiac contractile behavior, and opposing antiviral signatures support the use of gravitational unloading as a viable perturbation for studying mechanobiology in human tissue models. However, interpretation should remain bounded by the acute exposure, bulk-RNA-sequencing design and preliminary evidence for cellular causality. Additional context and validation can come from longer time courses, independent organoid lines, factorial interaction analyses, integration of in-flight astronaut data, cell-type-resolved measurements, and recovery experiments will be needed to determine whether the observed states represent transient adaptations or persistent dysfunction, and whether the transcriptomic-age shifts reflect sustained changes in tissue-aging trajectories.

## Methods

### Organoid culture

Five-month-old forebrain organoids from the F11362.1d1C11 iPSC line were obtained from the Neural Stem Cell Institute^17,18^. Neural organoids were cultured in non-treated 24-well plates (Cat# 10861-558, VWR) using Neurobasal-A medium (Cat# 10888-022, Gibco) supplemented with 2% B-27 (Cat# 12587010, Gibco), 2 mM GlutaMax (Cat# 35050-061, Gibco), and 1% penicillin/streptomycin (Cat# 15240-062, Gibco). Half the culture medium was replaced every two days. Cardiac organoids contained either iPSC-derived cardiomyocytes alone (CM) or cardiomyocytes cultured with endothelial cells (CMEC). Both preparations were maintained under 1g control conditions or exposed to simulated microgravity, as described below. Primary tonsil organoids were generated from dissociated human tonsil cells. For each organoid, 2-2.5 million cells were resuspended in 5 μL of complete medium consisting of RPMI 1640 with GlutaMax (Gibco, cat. no. 61870036), 10% fetal bovine serum (Corning, cat. no. 35010CV), 1% penicillin-streptomycin, 0.1 mM non-essential amino acids (Gibco, cat. no. 11140050), 1 mM sodium pyruvate (Gibco, cat. no. 11360070), 100 μg/ml Normocin (InvivoGen, cat. no. ant-nr), 1% Insulin-Transferrin-Selenium (Gibco, cat. no. 41400045), and 0.5 μg/ml human B-cell activating factor (BAFF) recombinant protein (Gibco, cat. no. PHC1674). The cell suspension was mixed with 10 μL of hydrogel, with or without 0.5 μL of LAIV vaccine per organoid. Following a 10-minute incubation period, each mixture was dropped into supporting medium (complete medium minus BAFF) in a 96-well plate to form tonsil organoids.

### Simulated microgravity

For simulated-microgravity exposure, 5–7 neural organoids or 3–4 tonsil organoids were placed in each 10-ml disposable high-aspect-ratio vessel (Synthecon, cat. no. D-410) and rotated for 24 hours at 25 or 28.3 rpm, respectively. Cardiac organoids from one well (containing 81 organoids) were loaded into each 2-ml autoclavable vessel (Synthecon) and rotated for 24 hours at 15 rpm. Parallel organoids maintained at 1g served as controls. After exposure, organoids were collected in either RNAlater Stabilization Solution (Invitrogen, cat. no. AM7020) or Allprotect Tissue Reagent (QIAGEN, cat. no. 76405) and stored at −80 °C.

### Cardiac motion analysis

Cardiac organoid motion was recorded at baseline and after 24 hours under 1g or simulated microgravity using 15-second grayscale videos (AVI format, 720p, 30 frames per second). Pixel-intensity changes across video frames were extracted using the ImageJ Time Series Analyzer macro (version 3.0) and analyzed using the R package gsignal.

### RNA sequencing and differential expression

Total RNA was extracted using the RNeasy Plus Micro Kit (Qiagen, cat. no. 74134) or AllPrep DNA/RNA/Protein Mini kit (Qiagen, cat. no. 80004), according to the manufacturer’s instructions. RNA quantification, library preparation, and mRNA sequencing were performed by Novogene (California, USA). Approximately 20 million paired-end, 150-bp read pairs were generated per sample on an Illumina NovaSeq 6000 Sequencing System. Raw FASTQ reads were trimmed using fastp^19^, aligned to the GRCh38 human reference genome using HISAT2^20^ and quantified with featureCounts^21^. Differential gene-expression analysis between groups was performed using the DESeq2 R package (version 1.38.3)^22^. Genes with adjusted P < 0.05 and |log2FC| > 0.58 were considered differentially expressed.

### Disease and gene-set enrichment

Differentially expressed genes were tested for overrepresentation of disease-associated gene sets from DisGeNET^13^ using clusterProfiler R package (version 4.6.2)^23^. Gene-set enrichment analysis against the C5 ontology gene-set collection was performed using GSEApy (version 1.0.5)^24^. Enrichment of gene sets representing the hallmarks of aging was analyzed using one-tailed, positive-enrichment gene-set analysis with fgsea (version 1.24.0)^25^.

### Rank-rank overlap analysis

Rank–rank hypergeometric overlap (RRHO) analysis compared ranked differential-expression lists restricted to genes shared by each pair of datasets. Overlap maps were generated using RRHO2 (version 1.0)^26^. Permutation P values were calculated using the pvalRRHO function in the RRHO R package (version 1.38.0)^27^. Comparisons included the organoid and published spaceflight signatures shown in Fig. 1e–g and the LAIV-associated and microgravity-associated signatures shown in Fig. 4c.

### Organoid-GTEx tissue correlations

GTEx RNA-sequencing read counts were obtained using the create_rse_manual function in recount3 (version 1.8.0)^28^. Read counts were converted to counts per million and log-transformed. For each tissue comparison, Spearman correlation coefficients were calculated using genes shared between the GTEx tissue samples and 1g organoid samples. Each GTEx tissue sample was compared with each organoid sample using the R stats package (R version 4.2.3).

### Transcriptomic age estimation

GTEx Analysis V8 RNA-sequencing counts, sample annotations and approved-access subject-age information were used to train tissue-specific transcriptomic age models. Counts were converted to counts per million and log-transformed. Lasso and elastic-net regularized generalized linear models were trained with 10-fold cross-validation using glmnet (version 4.1-8)^29^. The same expression preprocessing was applied to organoid samples, and the fitted models were used to estimate their transcriptomic ages under 1g and simulated-microgravity conditions. The original analysis specifies a one-tailed t-test for statistical comparison of the age estimates.

### Tonsil organoid transcriptomic age signature

Nominally significant up- and down-regulated genes were filtered using a P value cutoff of 0.001. GSVA was used to score each dataset using each gene set^30^. The up-regulated gene score was scaled from 0-1 to 20-70 to represent a “transcriptomic age” for the tonsil organoids.

## References

1. Tomsia, M. et al. Front. Physiol. 15, 1284644 (2024). doi 10.3389/fphys.2024.1284644

2. Roy-O’Reilly, M., Mulavara, A. & Williams, T. NPJ Microgravity 7, 5 (2021). doi 10.1038/s41526-021-00133-z

3. Vernice, N.A., Meydan, C., Afshinnekoo, E. & Mason, C.E. Precis. Clin. Med. 3, 284–291 (2020). doi 10.1093/pcmedi/pbaa022

4. Crucian, B.E. et al. Front. Immunol. 9, 1437 (2018). doi 10.3389/fimmu.2018.01437

5. Overbey, E.G. et al. Nature 632, 1145–1154 (2024). doi 10.1038/s41586-024-07639-y

6. da Silveira, W.A. et al. Cell 183, 1185–1201.e20 (2020). doi 10.1016/j.cell.2020.11.002

7. Hammond, T.G. & Hammond, J.M. Am. J. Physiol. Renal Physiol. 281, F12–F25 (2001). doi 10.1152/ajprenal.2001.281.1.F12

8. GTEx Consortium. Nat. Genet. 45, 580–585 (2013). doi 10.1038/ng.2653

9. Marotta, D. et al. Stem Cells Transl. Med. 13, 1186–1197 (2024). doi 10.1093/stcltm/szae070

10. Rampoldi, A. et al. Stem Cell Reports 17, 2272–2285 (2022). doi 10.1016/j.stemcr.2022.08.007

11. Wu, F. et al. Nat. Commun. 15, 4795 (2024). doi 10.1038/s41467-023-42013-y

12. López-Otín, C. et al. Cell 186, 243–278 (2023). doi 10.1016/j.cell.2022.11.001

13. Piñero, J. et al. Nucleic Acids Res. 45, D833–D839 (2017). doi 10.1093/nar/gkw943

14. Park, J.S., Davis, R.L. & Sue, C.M. Curr. Neurol. Neurosci. Rep. 18, 21 (2018). doi 10.1007/s11910-018-0829-3

15. Hayashi, J. & Carver, J.A. Biomolecules 12, 142 (2022). doi 10.3390/biom12010142

16. Wagar, L.E. et al. Nat. Med. 27, 125–135 (2021). doi 10.1038/s41591-020-01145-0

17. Karch, C.M. et al. Stem Cell Reports 13, 939–955 (2019). doi 10.1016/j.stemcr.2019.09.006

18. Bowles, K.R. et al. Cell 184, 4547–4563.e17 (2021). doi 10.1016/j.cell.2021.07.003

19. Chen, S., Zhou, Y., Chen, Y. & Gu, J. Bioinformatics 34, i884–i890 (2018). doi 10.1093/bioinformatics/bty560

20. Kim, D., Paggi, J.M., Park, C., Bennett, C. & Salzberg, S.L. Nat. Biotechnol. 37, 907–915 (2019). doi 10.1038/s41587-019-0201-4

21. Liao, Y., Smyth, G.K. & Shi, W. Bioinformatics 30, 923–930 (2014). doi 10.1093/bioinformatics/btt656

22. Love, M.I., Huber, W. & Anders, S. Genome Biol. 15, 550 (2014). doi 10.1186/s13059-014-0550-8

23. Yu, G., Wang, L.G., Han, Y. & He, Q.Y. OMICS 16, 284–287 (2012). doi 10.1089/omi.2011.0118

24. Fang, Z., Liu, X. & Peltz, G. Bioinformatics 39, btac757 (2023). doi 10.1093/bioinformatics/btac757

25. Korotkevich, G. et al. Preprint at bioRxiv (2016). doi 10.1101/060012

26. Cahill, K.M., Huo, Z., Tseng, G.C., Logan, R.W. & Seney, M.L. Sci. Rep. 8, 9588 (2018). doi 10.1038/s41598-018-27903-2

27. Plaisier, S.B., Taschereau, R., Wong, J.A. & Graeber, T.G. Nucleic Acids Res. 38, e169 (2010). doi 10.1093/nar/gkq636

28. Wilks, C. et al. Genome Biol. 22, 323 (2021). doi 10.1186/s13059-021-02533-6

29. Tay, J.K., Narasimhan, B. & Hastie, T. J. Stat. Softw. 106, 1–31 (2023). doi 10.18637/jss.v106.i01

30. Hänzelmann, S., Castelo, R. & Guinney, J. BMC Bioinformatics 14, 7 (2013). doi

